# Effects of Oral Exposure to HPAI H5N1 Pasteurized in Milk on Immune Response and Mortality in Mice

**DOI:** 10.1101/2024.10.03.616493

**Authors:** Pamela H. Brigleb, Ericka Kirkpatrick Roubidoux, Brandi Livingston, Bridgett Sharp, Victoria A. Meliopoulos, Shaoyuan Tan, Dorothea R. Morris, Lauren Lazure, Stacey Schultz-Cherry

## Abstract

In March 2024, there was the first reported outbreak of a highly pathogenic avian H5N1 influenza (HPAI) clade 2.3.4.4b virus in dairy cows in the United States. Since then, there have been several spillover events to cats, poultry, and humans. Multiple reports have discovered infectious virus in raw milk from infected dairy cows. Infectious virus can also last over a period on milking machine surfaces as a potential route of spread in cattle and contamination in raw milk. While the U.S. Food and Drug Administration has cleared commercial pasteurized milk as safe for consumption given the lack of infectious virus, there have been numerous reports that up to 30 percent of commercial milk tested were positive for HPAI H5N1 influenza virus genome copies. This is not necessarily unique to the HPAI H5N1 virus, as retrospective studies have identified H1N1 and H3N2 seropositivity in cows linked to decreased milk production. However, it is unknown how repeat exposure to the remaining viral proteins and genomic material in pasteurized milk modulates immune responses once ingested. We developed a successful in-house pasteurization protocol that inactivated high viral loads of the pandemic H1N1 strain A/California/04/2009 (Cal09) or bovine-derived HPAI H5N1 (A/bovine/Ohio.B24OSU-439/2024) viruses in raw milk. Mice were administered this milk daily for five days and rechallenged with each respective virus. We found that repeated oral exposure to inactivated virus was not sufficient to prevent or accelerate mortality from lethal challenge of HPAI H5N1, though it did result in a ∼0.5 log_10_ reduction viral titers in the brain and delayed clinical signs. In contrast, oral gavage of mice with pre-existing immunity to H1N1 influenza virus with virus pasteurized in milk were protected from morbidity and mortality upon bovine H5N1 viral challenge. These findings suggest that ingestion of inactivated HPAI H5N1 has limited potential health risk and does not prevent protective immune history-mediated responses to lethal infection.

## Introduction

Highly pathogenic avian influenza H5N1 viruses (HPAI) have been circulating in wild and domestic poultry populations since the early 2000s, resulting in spillover events into mammals, including humans (*1, 2*). In March 2024, HPAI H5N1 clade 2.3.4.4b was detected in dairy cows and cats in the United States (*3*–*5*). Since the first confirmed infection in March 2024, it has been detected in 14 states as of September 20, 2024 (*6*), with several case reports of spillover into humans, with clinical presentations including conjunctivitis and respiratory illness (*6*–*9*). With the average fatality of ∼50% in previously reported HPAI H5N1 human cases (*10*), it is crucial to understand the drivers of this outbreak in cattle and the potential risks it poses to humans.

One of the initial signs of dairy cattle infectivity was a decrease in feed intake, along with dropping milk production, a condition known as the ‘milk drop’ syndrome (*11, 12*). Upon further investigation, it was found that raw milk can contain high levels of HPAI H5N1 virus and that infectious virus remains infectious on the milking equipment, which may serve as a transmission route between cattle (*13*). This is not the first time influenza has been thought to infect cattle. H1N1 and H3N2 viruses have been previously linked to ‘milk drop’ syndrome and studies reported seroconversion by hemagglutination inhibition assays (HAI) and virus neutralization assays (*14, 15*).

While raw milk containing infectious virus can pose a threat to exposed workers and other animals or those who consume it, the US Food and Drug Administration maintains that commercial milk that is pasteurized is safe for human consumption. Several researchers have corroborated these findings with testing “in-house” pasteurization with either PCR machines or heat blocks as effective for inactivating infectious virus in contaminated milk, while some reported inconsistent results with the disclaimer that “in-house” pasteurization did not precisely replicate the industrial pasteurization process (*16*–*22*). For large batches, milk is typically pasteurized at 72°C for 15 seconds. Although the collective data demonstrates that there is no infectious HPAI H5N1 virus in commercial milk, several studies have reported the detection of viral genomic material in commercial milk products (*23*). However, there is limited research in understanding how the viral genomic material and any influenza viral proteins in pasteurized milk modulate immune responses, if any, and whether repeat exposure contributes to protection or accelerates mortality during lethal viral challenge. This study investigates the impact of repeat oral gavage of pasteurized H1N1 and a bovine-derived HPAI H5N1 A/bovine/Ohio.B24OSU-439/2024 clade 2.3.4.4b virus in milk on protection against lethal challenge. This study found that repeated oral exposure to inactivated HPAI H5N1 virus did not significantly affect mortality rates, although it reduced brain virus levels and delayed symptoms. However, mice that received oral gavage of milk-pasteurized virus and had prior immunity to H1N1 showed protection against severe outcomes from H5N1 infections. These studies highlight the safety of ingesting commercial milk positive for H5N1 virus genomic content and begin to uncover mechanisms of protection against lethal HPAI H5N1 viral infection.

## Results

### Successful heat inactivation of H1N1 and H5N1 viruses

To determine the effectiveness of heat inactivation following the guidelines for milk pasteurization, we conducted several inactivation studies in endpoint PCR or real-time PCR machines using the default settings with the lid temperature set to 105°C for A/California/04/2009 H1N1 (Cal09) or an endpoint PCR machine for A/bovine/Ohio.B24OSU-439/2024 H5N1 viruses. Previous studies have found that inactivation of H5N1 virus diluted in milk at 72°C for 15 seconds is sufficient, while others report some inconsistencies among repeats, with the caveat that in-house pasteurization does not fully recapitulate industrial-scale milk pasteurization. To test, PCR-confirmed influenza negative raw milk was obtained from a local dairy source and diluted 1:1 with H1N1 (1.58E+06 50% tissue-culture infectious dose (TCID_50_)) or H5N1 (1.58E+08 TCID_50_) virus and heated at 72°C for 15 seconds or 72°C for 30 seconds. Controls included a pasteurized milk-only control, to ensure the milk did not lyse the cells or eggs used for titer determination, and a sample where pasteurized milk was mixed with live virus 1:1 prior to cell or egg infectivity, to ensure that the milk does not contain anything that would inhibit viral replication, such as antiinfluenza antibodies. Samples were inoculated into Madin–Darby canine kidney (MDCK) cells and/or embryonated chicken eggs. Viral inactivation was assessed by TCID_50_ assay of which the limit of detection is (1.5 log_10_/ml) at 3 or 5 days-post inoculation (dpi) and/or by embryonated chicken eggs assessed for viral positivity by hemagglutination (HA) assay for increased viral replication sensitivity.

Using both endpoint and qPCR machines, we found that heat treatment of H1N1 (1.58 E+06 TCID_50_ total) and H5N1 viruses (1.58 E+08 TCID_50_ total) for 72°C for 15 seconds/30 seconds reduced viral titers to under the limit of detection (1.5 log_10_/ml) by TCID_50_ (Tables 1 and 2). After 3 days-post inoculation (dpi), all inoculated embryonated chicken eggs were negative by HA assay except for the H1N1 and H5N1 virus only controls (Tables 1 and 2). Pasteurized milk alone did not lyse MDCKs or eggs (Tables 1 and 2), and no significant difference was observed between the virus only controls and live virus added to milk, indicating that the milk itself does not inhibit viral replication (Tables 1 and 2) These results corroborate previous findings and highlight the importance of quality control checks for effective pasteurization for viral inactivation in both cells and eggs(*16*–*18, 21, 22, 24*). Milk pasteurized in-house at 72°C for 15 seconds, which also eliminated bacterial contaminants (Supplementary Figure 1) was used for the duration of the studies.

**Table 1.**
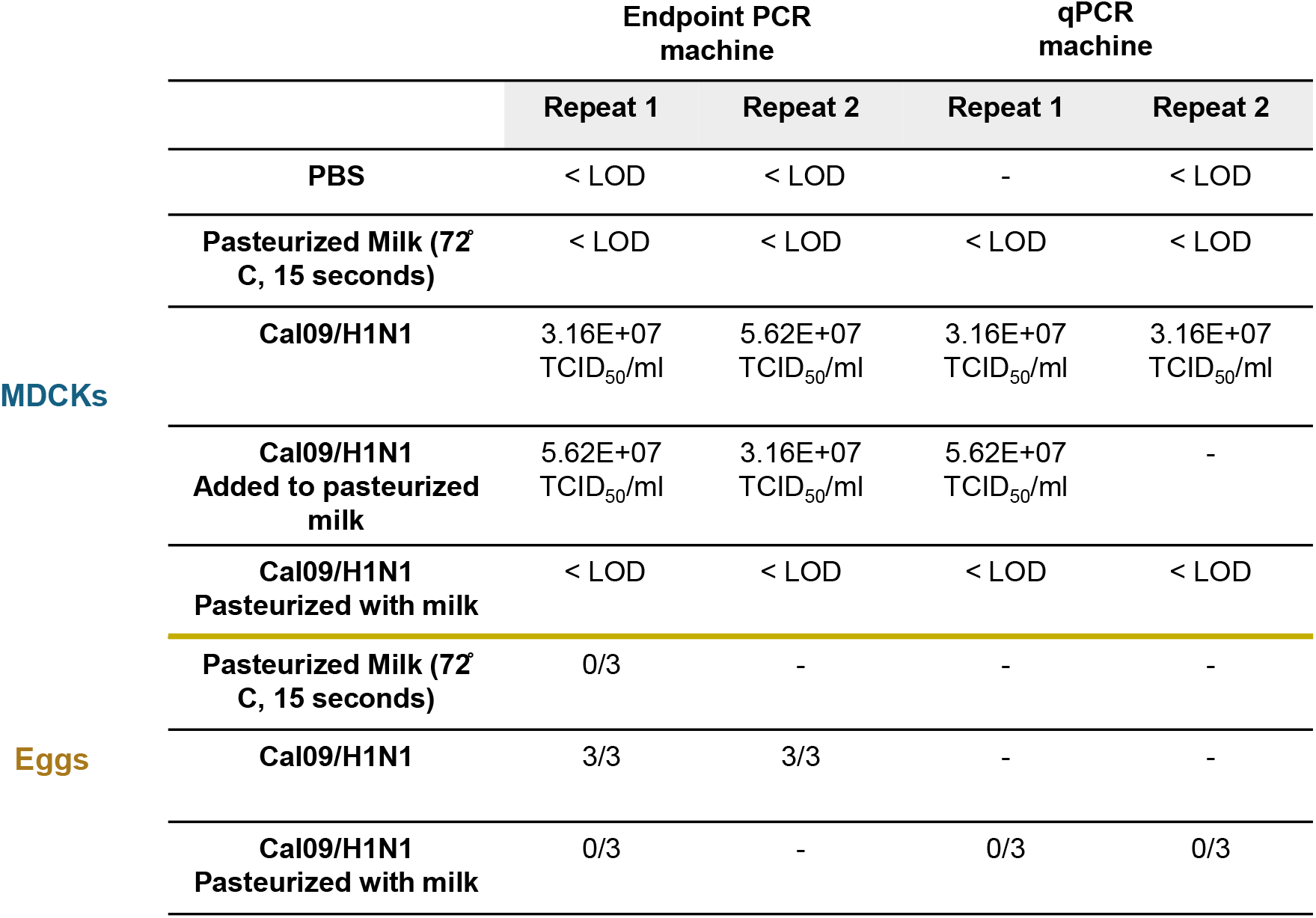
Characterization of milk pasteurization with H1N1 California 2009. Experiments were conducted with either an Endpoint PCR machine with lid closed or a real-time PCR machine with lid closed and set at the default settings of 105°C. Both PCR machines were programmed to a specific input volume to allow the internal temperature of the sample to reach 72°C prior to the program run. Samples were then either passaged through MDCKs or eggs at 3 days-post inoculation (dpi). Data is reported as a TCID_50_/ml, which was assessed using a hemagglutination assay and calculated with the Reed & Muench calculator, or by the number of eggs that were positive out of the total eggs infected. The limit of detection is 10^1.5^ TCID_50_/ml.

**Table 2.**
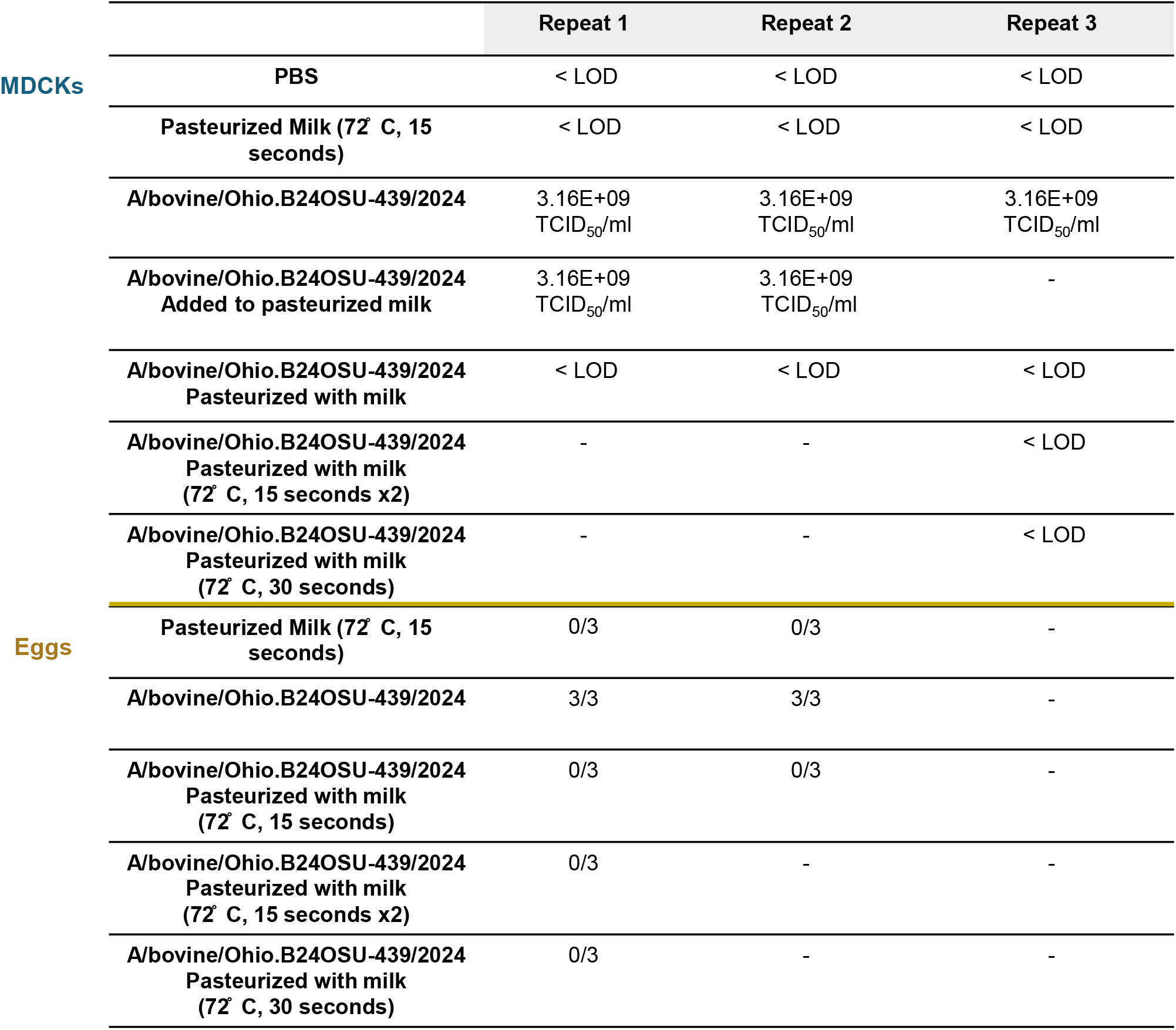
Characterization of pasteurization of raw milk and influenza H5N1 A/bovine/Ohio.B24OSU-439/2024. Experiments were conducted with an Endpoint PCR machine with lid closed, programmed to a specific input volume to allow the internal temperature of the sample to reach 72°C prior to the program run. Samples were then either passaged through MDCKs or eggs. Data is reported as a TCID_50_/ml, which was assessed using a hemagglutination assay and calculated with the Reed & Muench calculator, or by the number of eggs that were positive out of the total eggs infected at 36 h post-inoculation for H5N1 control virus or 3 days-post inoculation for the rest of the samples. The limit of detection is 10^1.5^ TCID_50_/ml.

### Impact of heat inactivated, pasteurization on viral protein stabilization

Viral genomic material has been detected in commercial milk supplies, with studies indicating that 20-30% of tested samples contain H5N1 genomic fragments (*23, 25, 26*). While the presence of these fragments does not pose a disease risk when consumed, the effects of the pasteurization process on viral proteins (as opposed to RNA) remain unclear. Thus, we investigated the stability of viral proteins following pasteurization in milk.

We observed viral protein degradation in samples heated by high-temperature pasteurization, 72°C for 15 seconds, that was even more pronounced in low-temperature pasteurization, 63°C for 30 minutes by total protein stain using Coomassie blue (Figure 1A). We next wanted to determine whether heat inactivation in milk would aid in the stabilization of viral proteins. We observed different viral protein degradation patterns when virus was pasteurized in PBS compared to milk following high-temperature pasteurization by total protein staining (Figure 1B). Interestingly, using HA-specific antibodies, we could detect HA in the virus pasteurized in milk, not in PBS, indicating that pasteurization of the virus in milk provides some protection against protein degradation (Figure 1C and 1D). We next investigated the impact of ingesting these viral proteins on protection from influenza infection.

**Figure 1.**
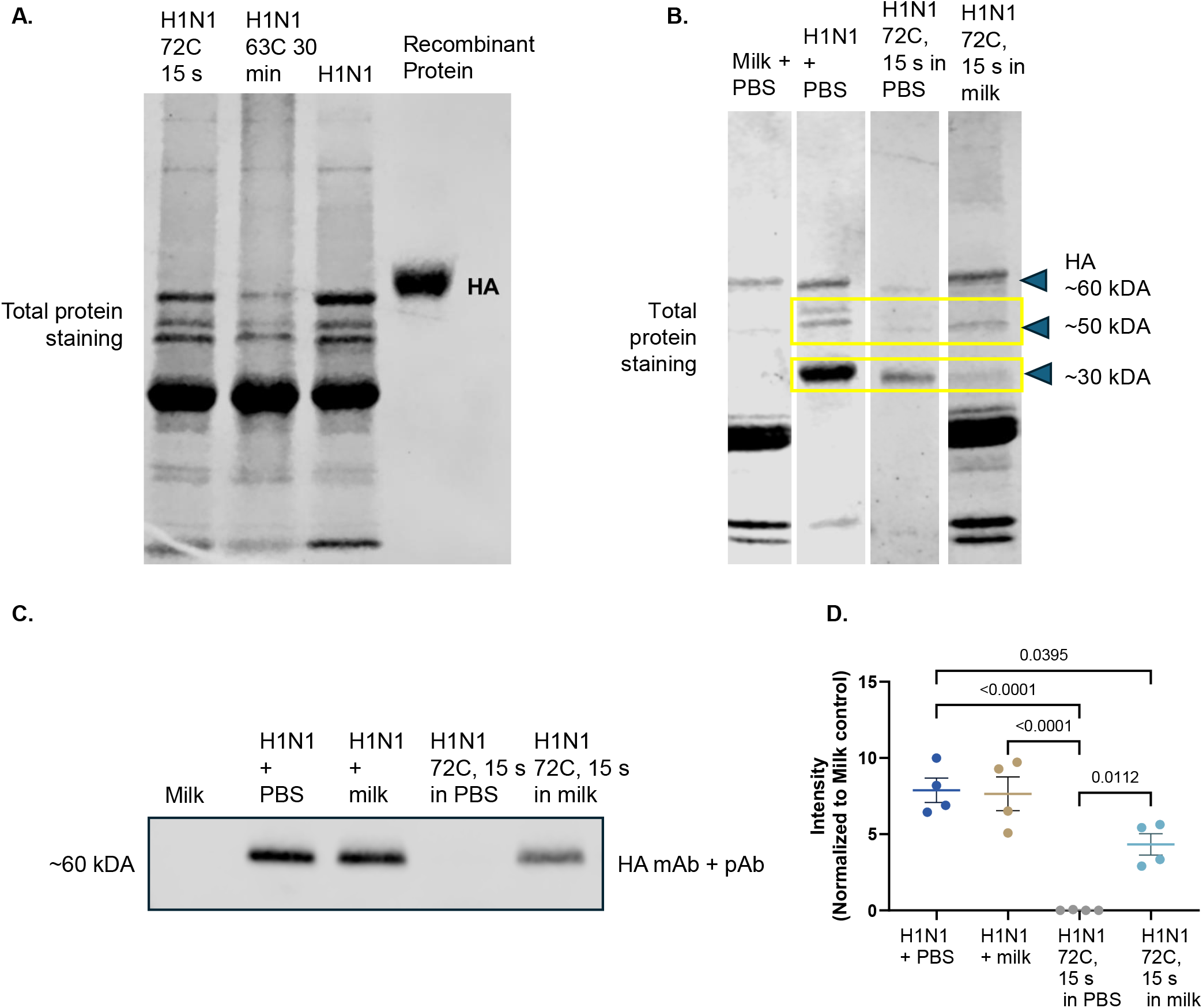
Pasteurization of H1N1 alters the composition of viral proteins. **(A)** Pasteurization of virus at either high temperature (72°C for 15 seconds) or low temperature (65°C for 30 minutes), H1N1 Cal09 or recombinant HA protein as a positive control was ran on a 4-20% gel and stained for total protein using Coomassie stain. **(B)** Total protein staining of diluted (1:50) of pasteurized milk in PBS, live H1N1 virus in PBS, or H1N1 virus pasteurized in either PBS or milk for 72°C for 15 seconds. Viral proteins with approximate molecular weight are shown in yellow boxes. **(B)** Combination of a monoclonal and polyclonal antibody against HA of pasteurized milk in PBS, live H1N1 virus in PBS, or H1N1 virus pasteurized in either PBS or milk for 72°C for 15 seconds. **(D)** Analysis of protein expression relative to milk in PBS control.

### Live, not inactivated, orally ingested H1N1 virus provides protection against lethal challenge

Previous studies have demonstrated that oral influenza vaccines may have some effectiveness (*27*–*30*). However, repeated protein exposure to a viral antigen could also lead to development of oral tolerance, thus hindering protective immune responses. Thus, we first asked how oral exposure to live or virus pasteurized in milk impacts response to lethal challenge using the pandemic Cal09 H1N1 virus. Mice were orally inoculated with pasteurized milk only, heat-inactivated H1N1 virus pasteurized in milk, or live virus diluted in milk daily for five days and monitored for morbidity (Figure 2A). The virus was diluted at a 1:1 ratio to milk, resulting in the mice receiving a 1.5×10^6^ TCID_50_ live or inactivated viral dose daily. All mice gained weight throughout the study and displayed no clinical signs or visible differences in fecal content, such as diarrhea, including the live virus group (Figure 2B). Twenty-one days after oral gavage, sera were collected to assess antibody titers and mice were intranasally inoculated with 5x the mean-lethal dose 50 (mLD_50_) of Cal09 H1N1 virus, including a naïve control group.

**Figure 2.**
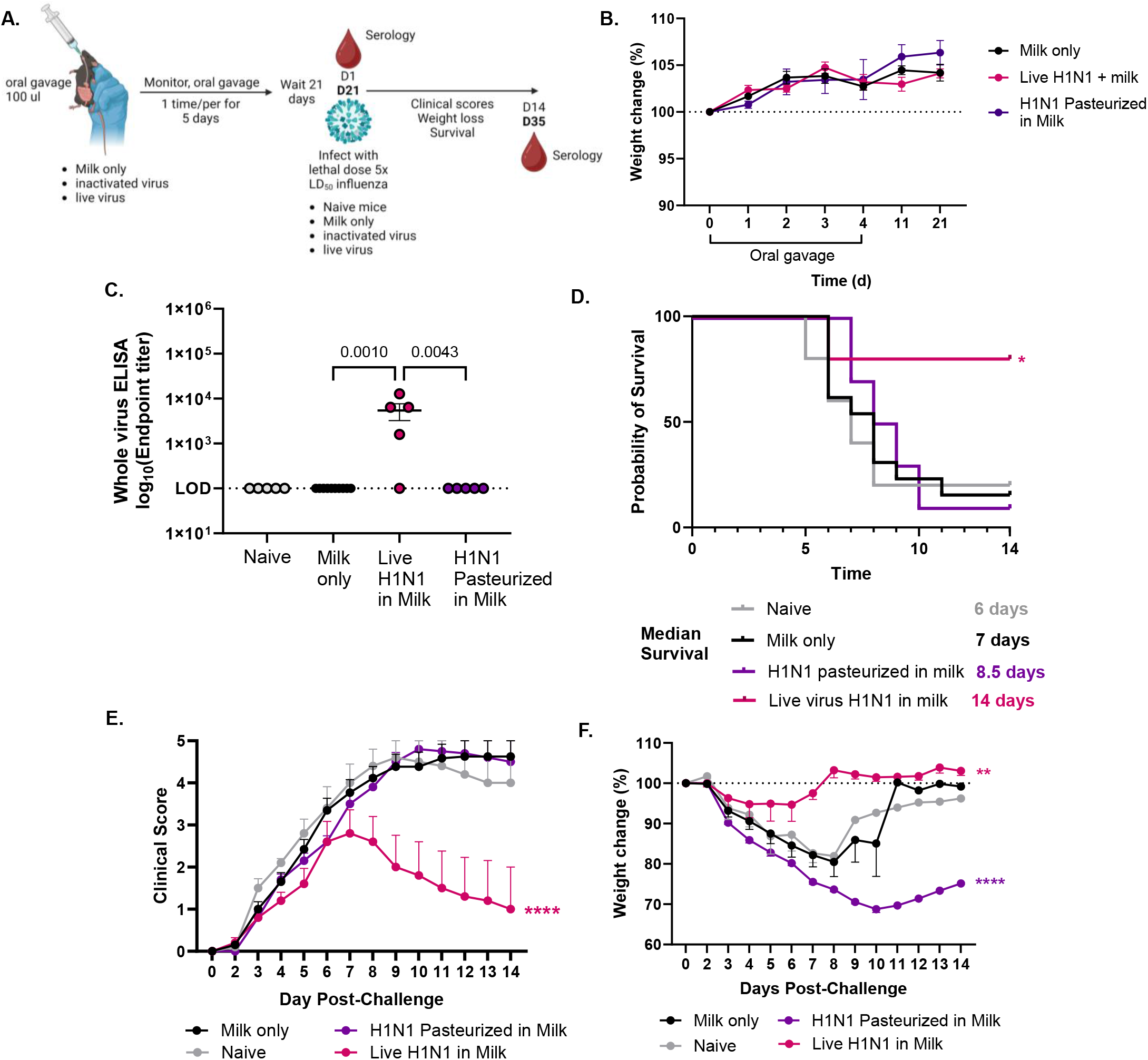
Live but not inactivated virus orally administered provides protection against lethal rechallenge to H5N1 in naïve WT mice. **(A)** Graphical summary of experimental design made in *Biorender*. **(B)** Weight change during and following oral gavage of pasteurized milk, 10^6^ live Cal09 H1N1, or virus pasteurized in milk. **(C)** Serum was collected from mice 21 days-post oral gavage start and prior to lethal challenge and antibodies against whole Cal09 H1N1 virus were assessed by ELISA. **(D-F)** Mice were rechallenged with 5x mLD_50_ 21 days-post start of oral gavage. **(D)** Survival **(E)** Clinical scores **(F)** Weight loss. n=5-15 mice/group with standard error of mean shown. Statistical analyses include Two-way ANOVA (B, D, E), Log-rank Mantel Cox test (C) and One-Way ANOVA with Tukey’s multiple comparisons test (F). * p<.05; **** p < .0001.

We found that oral gavage of live virus but not virus-pasteurized in milk induced systemic IgG antibodies (Figure 2C). In the live virus group, we observed significant protection from lethal challenge, with improved clinical scores and weight loss compared to the milk-only and naïve groups (Figures 2C-E). There were no differences between the milk-only and naïve groups, indicating that milk alone does not provide any protective responses against challenge (Figure 2C). Collectively, these results indicate that repeated oral ingestion of H1N1 virus inactivated in milk did not modulate responses to lethal infection, negatively or positively.

### Repeated ingestion of HPAI H5N1 pasteurized in milk does not negatively impact mortality or morbidity to lethal homologous viral challenge

To test the impact of orally digesting non-infectious H5N1 contaminated milk on subsequent exposure to H5N1 virus, mice were orally inoculated with pasteurized milk only or heat-inactivated virus diluted 1:1 in milk daily (1.5×10^8^ TCID_50_) for 5 days and monitored for morbidity (Figure 3). Mice continued to gain weight by 12 days-post gavage, similar to what we observed with the H1N1 virus (Figure 3B, Figure 2B). At 21 days-post start of oral gavage, sera were collected and mice were inoculated with either 10x or 1x the mLD_50_ of A/bovine/Ohio.B24OSU-439/2024 H5N1 virus, including a naïve control group. The mLD_50_ in male adult mice was 10 TCID50 (Supplementary Figure 2). At the high challenge dose of 10x mLD_50_, all mice succumbed or had to be humanely euthanized by 7 days post-inoculation (dpi) due to neurological presentation (Figure 3C). Despite the lack of protection against mortality, naïve and milk only groups displayed clinical signs prior to neurological symptoms, while the group that received H5N1 pasteurized in milk did not (Figure 3D). Furthermore, we observed ∼0.5 log10 reduction in brain viral titer at 7 dpi in the H5N1 pasteurized in milk (Figure 3E). However, we did not observe any differences in the lung titers amongst groups (Supplementary Figure 3). Due to the extreme lethality of this virus, we repeated this experiment with a lower lethal challenge of 1x the mLD_50_. We also found that repeated ingestion of H5N1 virus pasteurized in milk did not negatively impact mortality or morbidity to low-lethal challenge. Mortality was not significantly improved (Figure 3F) nor did we observe delayed clinical scores (Figure 3G). However, there was a mild reduction in brain but not lung titers in the H5N1 pasteurized in milk group, similar to the high-lethal dose (Figures 3H and Supplementary Figure 4). Collectively, these results suggest that repeated consumption of HPAI H5N1 virus pasteurized in milk, simulating viral genome positive commercial milk, does not adversely affect mortality or morbidity in response to a lethal homologous viral challenge.

**Figure 3.**
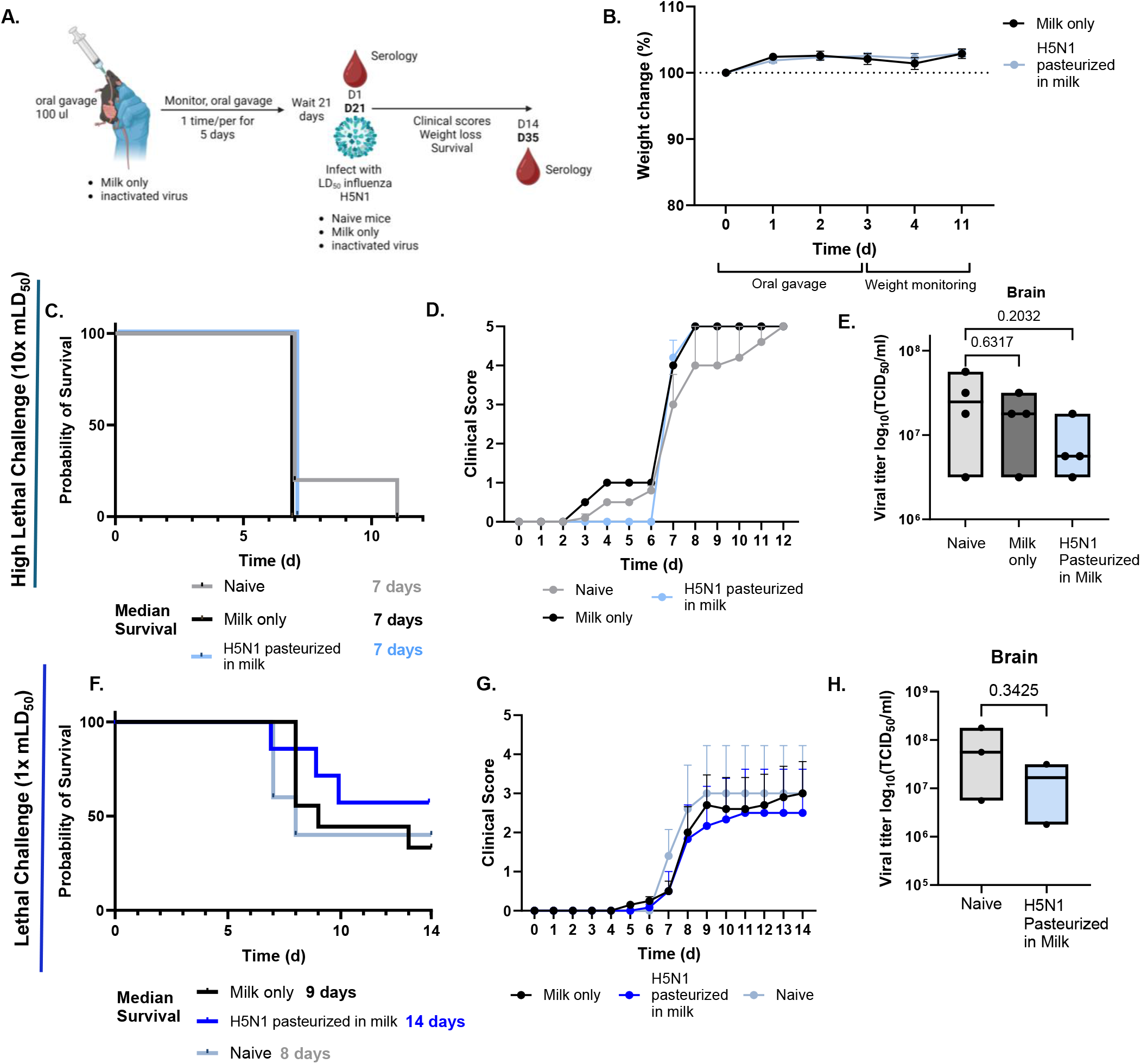
Repeated ingestion of HPAI H5N1 pasteurized in milk does not adversely affect mortality or morbidity in response to a lethal homologous viral challenge. **(A)** Graphical summary of experimental design made in *Biorender*. **(B)** Weight change during and following oral gavage of pasteurized milk or virus pasteurized in milk. **(C-E)** Mice were rechallenged with 10x mLD_50_ 21 days-post start of oral gavage. **(C)** Survival **(D)** Clinical scores **(E)** Brain titers. **(F-H)** Mice were rechallenged with 1x mLD_50_ 21 days-post start of oral gavage. **(F)** Survival **(G)** Clinical scores **(H)** Brain titers. n=4-9 mice/group with standard error of mean shown. Statistical analyses include One-Way ANOVA with Tukey’s multiple comparisons test (E, H).

**Figure 4.**
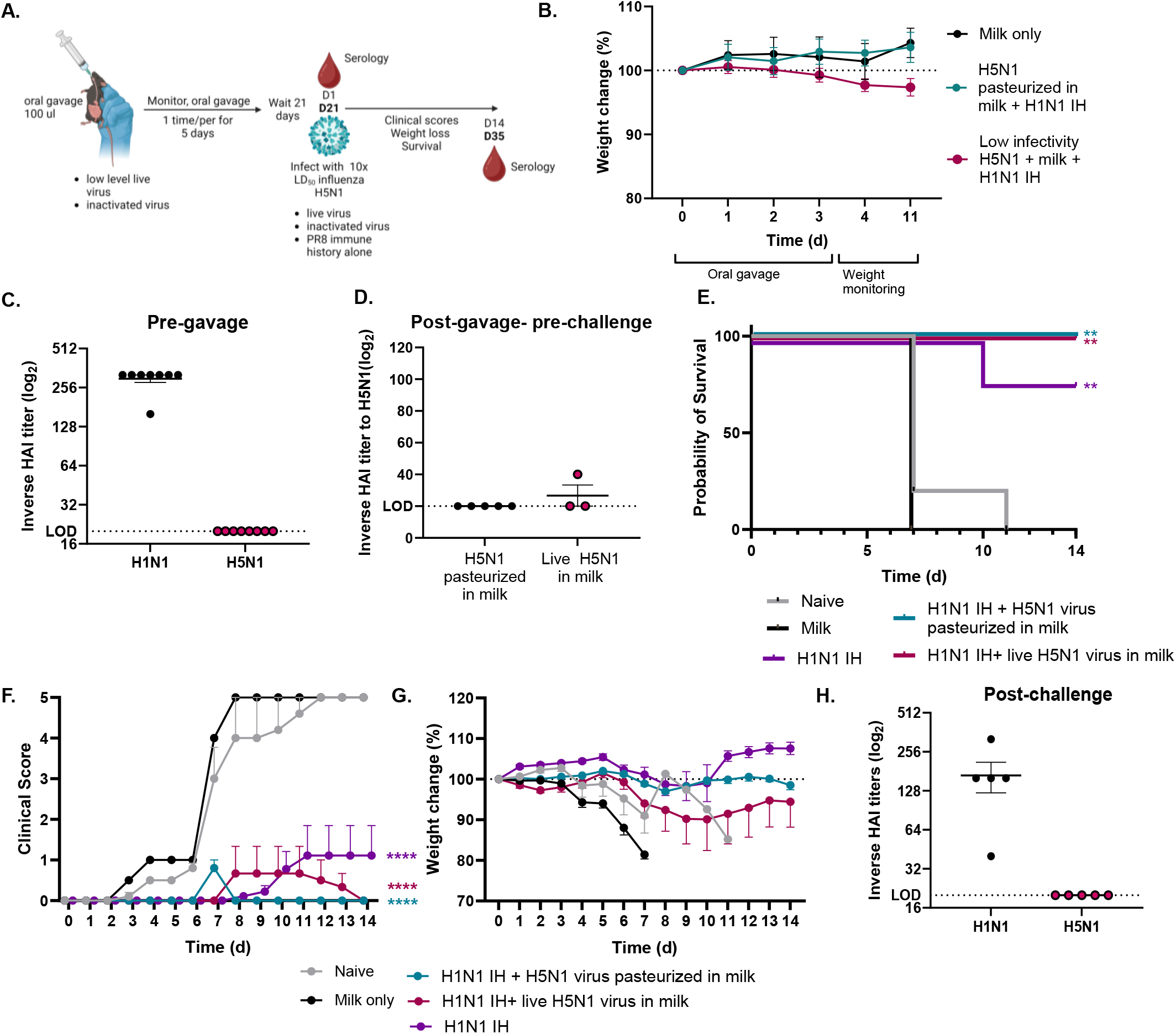
Prior H1N1 immune history provides protection against highly lethal HPAI H5N1 challenge and repeated ingestion of H5N1 contaminated pasteurized milk does not impede this protection. **(A)** Graphical summary of experimental design made in *Biorender*. **(B)** Weight change during and following oral gavage of pasteurized milk or virus pasteurized in milk. **(C, D)** sera was collected from mice prior to oral gavage, and 21 days-post oral gavage prior to lethal challenge and HAI titers were determined **(E-G)** Mice were rechallenged with 10x mLD_50_ 21 days-post start of oral gavage. **(E)** Survival **(F)** Clinical scores **(G)** Weight loss. **(H)** Serum was collected 21 days post-lethal challenge of survivors and HAI titers were determined. Note, these studies were conducted in parallel with Figure 3’s studies and the naïve and milk only control groups are the same between these two figures. n=3-9 mice/group with standard error of mean shown. Statistical analyses include Log-rank Mantel Cox test (E) and a Two-Way ANOVA (F, G) ** p<.01; **** p < .0001.

### H1N1 infection immune history provides protection against high lethal challenge of bovine H5N1

The studies described above were in naïve mice that had not previously seen influenza infection. However, most people have antibody responses to influenza virus from either vaccination or infection. To test if having immune history to H1N1 virus imposes any protection from subsequent H5N1 virus, C57BL/6J male mice were infected with A/Puerto Rico/8/1934 (PR8) H1N1 virus (2500-6500 EID_50_) followed by infection with A/Bovine/Ohio/2024 H5N1 3-14 weeks later or gavaged with either milk, H5N1 virus pasteurized in milk, or low infectious virus in pasteurized milk (Figure 4A). Mice with H1N1 virus infection immune history were protected from mortality and had minimal weight loss, whereas weight loss was not observed in milk only and H5N1 virus pasteurized in milk groups (Figure 4B). These results indicate that prior immune history with an H1N1 strain provides protection against oral ingestion of lowly-positive milk.

We next investigated (i) whether infection immune history alone from a genetically different H1N1 virus provides protection or (ii) whether repeated oral ingestion of H5N1 virus pasteurized in milk would boost or inhibit prior immune history responses against high lethal H5N1 challenge (10x mLD_50_). Since these studies were conducted in parallel, the naïve and milk only control groups are the same as reported previously (Figure 3). We found that mice that had H1N1 immune history with repeated oral ingestion of live or pasteurized H5N1 virus were 100% protected from high lethal challenge of H5N1 compared to 80% protective with immune history alone (Figure 4C). Mild clinical scores and weight loss was observed in the group that received repeated oral ingestion of H5N1 virus pasteurized in milk (Figure 4D). These results were interesting, as the mice with H1N1 immune history had high H1N1 HAI titers but little-to-no H5N1 virus HAI titers pre-challenge, pre-gavage, and post-challenge (Figures 4C, D, & H). Amino acid alignment demonstrated that A/Bovine/Ohio/24 H5N1 and A/PR8 H1N1 viruses have a 64.34% HA homology, but higher for NA (83.58%) and NP (93.98%) (Supplementary Figure 4). These data suggest that there is an alternative correlate of protection, independent of HA neutralizing antibodies, that warrants further investigation. Collectively, these results suggest that H1N1 prior immunity provides protection against lethal bovine H5N1 challenge and that ingestion of HPAI contaminated commercial milk does not impede the protective immune responses facilitated by infection H1N1 immune history.

## Discussion

Since March 2024, there has been an escalation in the reports of HPAI H5N1 clade 2.3.4.4b influenza virus infected cattle and crossover to other mammals, including humans (*3*). Previous data has implicated that other influenza A strains, including H1N1 and H3N2, can infect cattle resulting in “milk drop” syndrome (*14, 15*). Understanding the mechanisms that allow cross-mammalian infectivity of HPAI H5N1 virus is essential. Furthermore, with high levels of H5N1 genomic contents being detected in commercial milk, it is also essential to understand the potential risks of repeatedly consuming contaminated, but not infectious, HPAI milk on our immune responses.

Ingesting high levels of viral proteins, even those inactivated, can modulate immune responses. This concept is the basis for developing oral vaccines, of which there were several in the pipeline for influenza (*27*–*30*). It is how livestock, including chickens, are immunized to several diseases (*31*). In theory, ingesting inactivated viral proteins can result in protective immune responses based on developing antibodies against those proteins or promote oral tolerance, which could inhibit inflammatory responses to the viral protein and may accelerate mortality or morbidity to the pathogen. We demonstrated that heat inactivation degraded viral proteins, but inactivation at 72°C for 15 seconds in milk, not PBS, still allowed the binding of a combination of monoclonal and polyclonal antibodies against HA to the protein, indicating that virus pasteurization in milk may aid viral protein stability. While high levels of HPAI H5N1 genomic content can be detected in commercial milk, it is unknown to the extent to which viral proteins, degraded or not degraded, are present. Since it is likely, based on our studies, that antibody-based detection for all viral proteins might not work due to protein degradation, assays such as mass spectrometry might be necessary to determine viral protein presence in commercial milk. Although, since there is infectious virus in raw milk that is very stable over time, there is highly likely some level of viral proteins also in commercial milk.

This study aimed to do a proof-of-concept, where controlled but high amounts of H1N1 or H5N1 were added to raw milk and then pasteurized at 72°C for 15 seconds. We chose this experimental design to accurately control how much virus or inactivated virus the mice were administered daily. We did so at the highest dose possible, for if we did not observe any impact on challenge responses at this high dose, we would not expect to observe differences with either commercial milk positive for H5N1 genomic content or lower doses of spiked-in virus. Based on the inconsistent results from previous papers with in-house pasteurization, we tested our pasteurization technique multiple ways: standard MDCK-based TCID_50_ for three days, an extended TCID_50_ for five days, and in eggs. Based on these results under our specific conditions, we observed repeated successful inactivation of high-titer H1N1 and H5N1. However, if high titers of H5N1 or other influenza strains are detected in raw milk, it is essential for routine screening in both cell and egg-based assays to ensure proper inactivation, as 3/5 mice died from accidental administration of insufficiently inactivated H5N1, most likely due to human error (data not shown) similar to prior findings (*32*).

Repeat oral gavage of high-titer inactivated H5N1 virus had mild-to-low protection against high lethal challenge, including that no clinical scores were reported until the virus went neurologic at day 7, and a slight reduction (mean was ∼0.5 log less compared to naïve control) of viral titer in the brain. This bovine isolate of HPAI H5N1 is exceptionally lethal, with the mLD_50_ of 10^1^ TCID_50_. Interestingly, we showed that prior infection history with PR8 H1N1 virus was sufficient to protect against orally ingesting contaminated H5N1 milk. Furthermore, PR8 H1N1 virus immune history mice who were orally gavaged with inactivated H5N1 virus were 100% protected against mortality with mild signs of morbidity to high titer challenge of HPAI H5N1 (10x mLD_50_). However, we detected little-to-no H5N1 HAI titers in the immune history mice that were protected, even following a 10x mLD_50_ challenge. PR8 and our strain of bovine H5N1 virus only have a 64.34% similarity in the HA and an 83.58% similarity in the NA amino acid sequences. Previous research has speculated that pandemic and other H1N1 strains can provide immunity to H5N1 viruses in an NA-dependent manner (*33*), which may be also be important in this model. These data collectively suggest that PR8 infection history protects against lethal HPAI H5N1 challenge which is not impeded when repeatedly exposed to H5N1 virus contaminated pasteurized milk.

There are several limitations to this study. Due to the urgency of this research with the HPAI H5N1 outbreak, we used accessible mice, especially for immune history studies and we focused these studies on solely male, adult C57BL/6J mice. These findings may differ in female mice or different mouse strains. Furthermore, since we repurposed mice with prior PR8 infection history, there was a limitation in the sample size for the PR8 immune history with live or inactivated milk. There was a range of immune history from 2500-6500 EID_50_ of PR8 virus, although antibody responses to PR8 and H5N1 viruses were assessed pre-oral gavage and pre-challenge to control for this variable.

While we have known about viral genomic content being detected in commercial milk, it was unclear what the impact of this genomic content and residual viral protein on immune responses. We aimed to assess this question using two different influenza A virus strains and added the complexity of whether immune history to a very different influenza A virus would also play a role. While there is still a lot of research to be done in this field to answer a complex question, including the testing of H5N1 PCR positive commercial milk and whether repeated exposure to that would modulate immune responses, and different types of immune history matter, we emphasize the need for further research in this area. These studies emphasize the safety of consuming commercial milk that contains H5N1 virus proteomic material and begin to reveal the mechanisms that provide protection against lethal HPAI H5N1 viral infections.

## Materials and Methods

### Animal husbandry and ethics statement

All procedures were approved by the Institutional Biosafety Committee and the Animal Care and Use Committee at St. Jude Children’s Research Hospital in compliance with the Guide for the Care and Use of Laboratory Animals. These guidelines were developed by the Institute of Laboratory Animal Resources and approved by the Governing Board of the U.S. National Research Council. Mice were kept under 12 h light/dark cycles at an ambient temperature of 68 °F and 45% humidity. They had continuous access to diet and water. When required, either for humane endpoints or timepoint collection, mice were euthanized following American Veterinary Medical Association guidelines.

### Biosafety

All experiments using the highly pathogenic avian influenza strain were conducted in a biosafety level 3 enhanced containment laboratory. Investigators were required to wear appropriate respirator equipment (RACAL, Health and Safety Inc., Frederick, MD) and PPE. Mice were housed in negative pressure, HEPA-filtered and vented isolation containers. Experiments using the pandemic California 2009 H1N1 were conducted in a biosafety level 2.

### Cells and viruses

Madin-Darby canine kidney (MDCK, American Type Culture Collection, CCL-34) cells were cultured in Dulbecco’s minimum essential medium (DMEM; Corning) supplemented with 2 mM GlutaMAX (Gibco) and 10% fetal bovine serum (FBS; HyClone) and grown at 37°C with 5% CO2. A/California/04/2009 (H1N1) viruses and A/bovine/Ohio.B24OSU-439/2024 were kind gifts from Robert Webster and the Webby lab at St. Jude Children’s Research Hospital, respectively. The viruses were propagated in the allantoic cavity of 10-day-old specific-pathogen-free embryonated chicken eggs at 37°C as previously described (74). Briefly, allantoic fluid was cleared by centrifugation following harvest and then stored at −80°C (13, 14). Viral titers were determined by 50% tissue culture infective dose (TCID50) analysis (see below).

### Pasteurization

Raw cows’ milk was used in these studies that was tested by qPCR to be negative for both H5N1 and influenza A (data not shown). Milk with or without virus was pasteurized in PCR tubes with a maximum volume of 50 μL in either an endpoint or real-time PCR machine, as labeled in the figures. Pasteurization occurred for either 72°C for 15 seconds, 72°C for 30 seconds, or 63°C for 30 minutes.

### Oral inoculation of mice

Ten-to-sixteen-week-old male C57BL/6J mice (Jackson Laboratories) were perorally inoculated with sterile disposable plastic oral gavage needles once a day for a total of five days. Mice were orally gavage with 100 μL of pasteurized milk diluted in phosphate-buffered saline (PBS), virus (H1N1 Cal09 or H5N1 Ohio24) pasteurized in milk, or live virus diluted in pasteurized milk. Mice were monitored and weighed daily at time of gavage and monitored daily after up to 7 days total and then weekly. Fecal contents were also monitored for diarrhea, but no mice displayed signs of intestinal upset or lost weight following oral gavage with pasteurized milk alone.

### Mouse lethal dose 50 determination

Ten-to-twelve-week-old male C57BL/6J were lightly anaesthetized with inhaled isoflurane and intranasally (IN) inoculated with 10^1^, 10^2^, or 10^3^ TCID_50_ of Ohio24 H5N1 diluted in PBS for a total volume of 25 μL (2-4 mice per dosage). Body weight, clinical scores, and survival were monitored daily for 13 days. Mice were humanly euthanized if they lost more than 30% body weight and/or displayed clinical scores of 3 or above. These included neurological symptoms, which were observed in most euthanized mice, including circling, tremors, and laying on one side with limited motility.

### Viral titer determination

Viral titers were determined as previously described by 50% tissue culture infectious dose (TCID_50_) assays. Briefly, confluent MDCK cells were infected in duplicate or triplicate with 10-fold dilutions pasteurization samples or tissue homogenates (bead beat in 500 μl PBS) in 100 μl of minimal essential medium (MEM) plus 0.75% bovine serum albumin (BSA) and 1 μg/ml tosylsulfonyl phenylalanyl chloromethyl ketone (TPCK)-treated trypsin (H1N1 samples only). After 3-5 days of incubation at 37 °C and 5% CO2, 50 μl of the supernatant was combined and mixed with 50 μl of 0.5% packed turkey red blood cells diluted in PBS for 45 minutes at room temperature and scored by hemagglutination (HA) endpoint. Infectious viral titers were calculated using the Reed-Muench method (*34*).

### Mouse viral infectivity

At timepoints outlined per figure, mice were challenged by IN inoculation with either 10x or 1x mLD_50_, as determined in this or previous studies, with A/bovine/Ohio.B24OSU-439/2024 or 5x the mLD_50_ for A/California/04/2009 diluted in PBS to a total volume of 25 μL that were lightly anesthetized with inhaled isoflurane. Body weight, clinical scores, and survival were monitored daily for 14 days. Moribund mice that lost more than 30% body weight and/or reached clinical scores of greater than four were humanely euthanized. Clinical signs were scored as follows: 0, no observable signs; 1, active, squinting, mild hunch or scruffy appearance; 2, squinting and hunching, ruffled fur, but active; 3, excessive hunching, squinting, visible weight loss, not active when stimulated; 4, not active when stimulated, sunken face and severe weight loss, shivering, rapid breathing, moribund; and 5, death. For mice infected with H5N1, the majority also displayed neurological signs, including circling, tremors, and laying on one side with limited motility. Mice were humanly euthanized when these neurological signs were present. Post-euthanasia, the whole lung and brain were harvested and stored at −80 °C for tissue processing and future analysis.

### Viral growth in embryonated chicken eggs

Ten-day-old embryonated chicken eggs were inoculated with samples described in Tables 1 and 2. All samples were diluted at a 1:2 except for the virus only positive controls and conducted in triplicate. Egg sampling occurred daily with a small aliquot of allantoic fluid and assessed for viral positivity by HA assay as previously described (add reference). At 36 hours-post inoculation, the positive control H5N1-infected eggs were killed by incubation overnight at 4°C. The remainder of the eggs inoculated with other samples were allowed to grow for 3 days, at which time the eggs were killed by incubation overnight at 4°C.

### Immunoblots

Samples were processed and loaded on a sodium dodecyl sulfate-polyacrylamide gel electrophoresis (4%–20% tris-glycine 1.0 mm Mini Protein Gels from Invitrogen). The gel was either stained with Coomassie blue and de-stained overnight prior to imaging, or was transferred to a nitrocellulose membrane using iBlock transfer stacks (ThermoFisher IB24002). They were then probed for protein with a combination of anti-HA monoclonal and polyclonal Abs (Sino Biologicals 11055-RM10 and 11055-T62). IRDye 680RD goat anti-rabbit IgG secondary (LI-COR 926-68071) antibody using the iBind (ThermoFisher) according to the manufacturer’s instructions. The blot was imaged on the LI-COR Odyssey.

### Bacterial screening

Raw milk was pasteurized using the methods described above. In triplicate, raw milk only or raw milk that had been pasteurized were spot plated following serial dilutions on blood agar plates and placed in either aerobic or anaerobic chambers overnight at 37°C. No difference was detected in the colony size or number between the aerobic or anaerobic growth conditions for the raw milk. The number of colonies was enumerated and the Colony forming units (CFU) per ml was calculated. No bacterial colonies were detected in either condition in the in-house pasteurized milk.

### Antibody quantification

Whole virus (H1N1 or H5N1) enzyme linked immunosorbent assays (ELISAs) were conducted using 384-well flat-bottom MaxiSorp plates (ThermoFisher) coated with either purified A/California/04/2009 at 5μg/ml or 10^6^ TCID_50_ of the virus stock of A/bovine/Ohio.B24OSU-439/2024 overnight at 4 °C. Plates were washed 4 times with PBS containing 0.1% Tween-20 (PBS-T) using the AquaMax 4000 plate washer system for H1N1 ELISA’s or handwashed for H5N1 ELISAs. Plates were blocked with PBS-T containing 0.5% Omniblok non-fat milk powder (AmericanBio) and 3% goat serum (Gibco) for 1 h at room temperature. The wash buffer was removed, and plates were tapped dry. Mouse sera was diluted 1:5 in PBS and ran in duplicate. Positive and negative mouse sera was used as controls for both sets of ELISAs. Samples were incubated at room temperature for 2 hours and then washed 4 times with PBS-T. Anti-mouse peroxidase-conjugated IgG secondary antibody was diluted at 1:3000 (Invitrogen 62-6520) in blocking buffer and 15 μL was added per well and incubated at room temperature for 1 hour. Plates were washed 4 times with PBS-T and developed using SIGMAFAST™ OPD (Sigma-Aldrich) for 10 min at room temperature. Plates were read at 490 nm using a BioTek Synergy2 plate reader and Gen5 (v3.09) software. For each plate, an upper 99% confidence interval (CI) of blank wells OD values was determined and used in determining the endpoint titers. Alternatively, mouse sera were treated with receptor-destroying enzyme (RDE; Seiken 370013), and HAI assays were performed as described (*35, 36*).

### Immune History

Adult male WT C57BL/6J mice were inoculated by IN with 2500, 4500, or 6500 Egg infective dose at 50% (EID_50_) of A/Puerto Rico/8/1934 (H1N1) and monitored daily for 14 days post-infection. A minimum of 3 weeks post-infection, mice were bled via eye bleeds prior to oral gavage to determine antibody titers or prior to challenge with lethal A/bovine/Ohio.B24OSU-439/2024.

### Phylogenetic Tree Construction

H5N1 sequences were obtained from the EpiFlu database of the Global Initiative on Sharing All Influenza Data (GISAID) (https://doi.org/10.2807/1560-7917.ES.2017.22.13.30494). We downloaded all complete sequences from human, mammalian, and avian hosts in clade 2.3.4.4b from North and South America, yielding a total of 6,480 strains as of August 23, 2024. To select optimal representative sequences for phylogenetic analysis, an initial phylogenetic tree was constructed using segment HA from all downloaded strains. The large phylogeny was then down-sampled using PARNAS(*37*). The optimal representative sequences, along with our lab strain A/bovine/Ohio/B24OSU-439/2024, were aligned using MUSCLE (*38*) and the sequence ends were trimmed to equal lengths in AliView manually (*39*). Maximum likelihood trees were then constructed using IQ-TREE2 with the GTR+F+R5 model and 1,000 bootstrap replicates (*40*). The resulting tree files were visualized using Figtree (http://tree.bio.ed.ac.uk/software/figtree/) (41).

### Pairwise Alignment, Identity, and Mutation Detection

Pairwise amino acid sequence alignments were performed in R using the pairwiseAlignment function from the pwalign package, and sequence identity was calculated using the pid function: (https://bioconductor.org/packages/pwalign).

### Statistical analyses

All animals were randomly selected for each control and experimental group. Male mice were selected due to current mouse availability, but all experiments should be repeated with female mice to determine whether there are any sex biases in the data. A limitation to this study was low sample size for some experiments, due to mouse availability (immune history mice), the urgency of these studies and limited access to our BSL3 facility. All graphs and statistical analyses were conducted using GraphPad Prism version 10.0. and described in each figure legend if applicable. The sample size and number of repeats is reported per figure. Statistical analyses include a One-way ANOVA with Tukey’s Multiple Comparisons test, Two-way ANOVA (mixed model) with multiple comparisons, and a log-rank Mantel-Cox test for survival curves and is also reported in the figure legend.

## Acknowledgements

This project has been funded in whole or in part with Federal funds from the National Institute of Allergy and Infectious Diseases, National Institutes of Health, Department of Health and Human Services, under Contract No. 75N93021C00016 (S.S.-C.), the American Lebanese Syrian Associated Charities (ALSAC) (S.S.-C.), by the National Institute of Health, Institutional Postdoctoral Training Grants (T32) Infectious Disease Therapeutics T32AI106700-08 (P.H.B.), and the National Institute of Health, Ruth L. Kirschstein Postdoctoral Individual National Research Service Award F32AI183804 (P.H.B).

The authors would like to extend their gratitude toward Richard Webby and the Webby lab at St. Jude Children’s Research Hospital for providing the original bovine isolate of HPAI H5N1 (A/bovine/Ohio.B24OSU-439/2024). In addition, the authors would like to thank Benjamin Treat and the Margolis Lab (Elisa Margolis) for providing Columbia blood agar plates and plating the bacteria samples for the raw and pasteurized milk samples. We would also like to extend our gratitude to the St. Jude Children’s Research Hospital Animal Facility staff and those who operate the Biosafety Level 3 facility. Illustrations in Figures 2-4 created with BioRender.com

## Author contributions

Conceptualization: S.S.-C., P.H.B., E.K.R., Methodology: S.S.-C.P.H.B., E.K.R., B.L., B.S., S.T. Investigation: S.S.-C., P.H.B., E.K.R., B.L., B.S., D.R.M., V.A.M., L.L., S.T. Visualization: S.S.-C., P.H.B., E.K.R., S.T. Funding acquisition: S.S.-C., P.H.B. Project administration: S.S.-C., P.H.B., E.K.R. Supervision: S.S.-C., P.H.B., E.K.R. Writing – original draft: S.S.-C., P.H.B., E.K.R. Writing – review & editing: S.S.-C., P.H.B., E.K.R., S.T., B.S.

## Competing interests

Authors declare that they have no competing interests.

## Data availability

All data are available in the main text or the supplementary materials.

## Notes

### Competing Interest Statement

The authors have declared no competing interest.

