## Supplemental Material for "Effects of Oral Exposure to HPAI H5N1 Pasteurized in Milk on Immune Response and Mortality in Mice"

A.

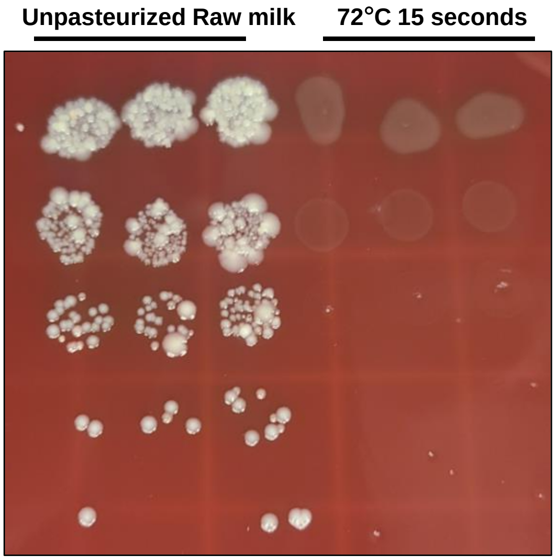


| | Unpasteurized milk | 72C, 15 seconds pasteurization |
|---|---|---|
| **Bacteria concentration** | 3.1 x 10^5 CFU/ml | < LOD |

**Supplementary Figure 1.** Pasteurization protocol successfully kills bacteria in raw milk. Raw milk or milk pasteurized using a real-time PCR machine for 72°C for 15 seconds was serially diluted on a blood agar plate in triplicate and allowed to grow overnight at 37°C.

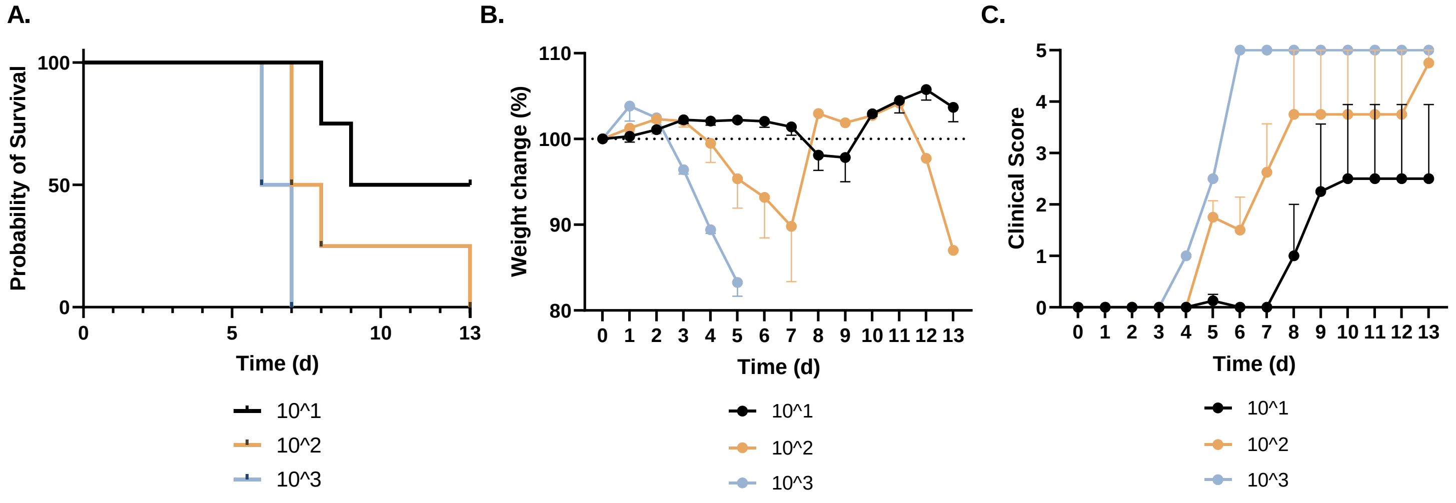


**Supplementary Figure 2.** $LD_{50}$ in adult male WT C57Bl/6J male mice. Mice were intranasally inoculated with $10^1$ (n=4), $10^2$ (n=4) or $10^3$ (n=2) $TCID_{50}$ of A/bovine/Ohio.B24OSU-439/2024 H5N1. Mice were weighed and monitored daily, and the majority succumbed to infection displaying neurological symptoms. The $LD_{50}$ in adult male WT C57Bl/6J male mice was determined as $10^1$ $TCID_{50,}$ and this dose was used for challenge studies with H5N1 marked as $LD_{50}$ challenge.

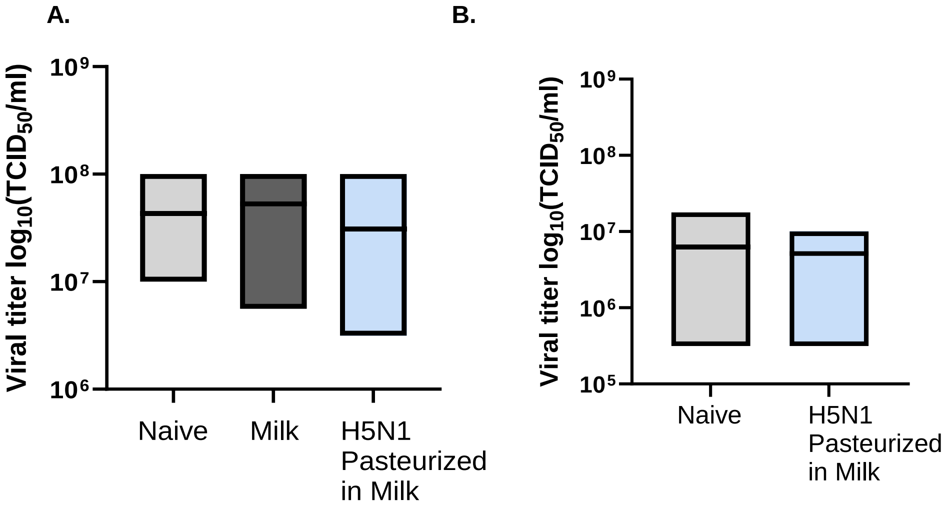


**Supplementary Figure 3. Repeated ingestion of HPAI H5N1 pasteurized in milk does not impact lung titers following H5N1 challenge.** **(A)** Graphical summary of experimental design made in *Biorender.* **(B)** Weight change during and following oral gavage of pasteurized milk or virus pasteurized in milk. **(C-E)** Mice were rechallenged with 10x $mLD_{50}$ 21 days-post start of oral gavage. **(C)** Survival **(D)** Clinical scores **(E)** Brain titers. **(F-H)** Mice were rechallenged with 1x $mLD_{50}$ 21 days-post start of oral gavage. **(F)** Survival **(G)** Clinical scores **(H)** Brain titers. n=4-9 mice/group with standard error of mean shown. Statistical analyses include One-Way ANOVA with Tukey's multiple comparisons test (E, H).

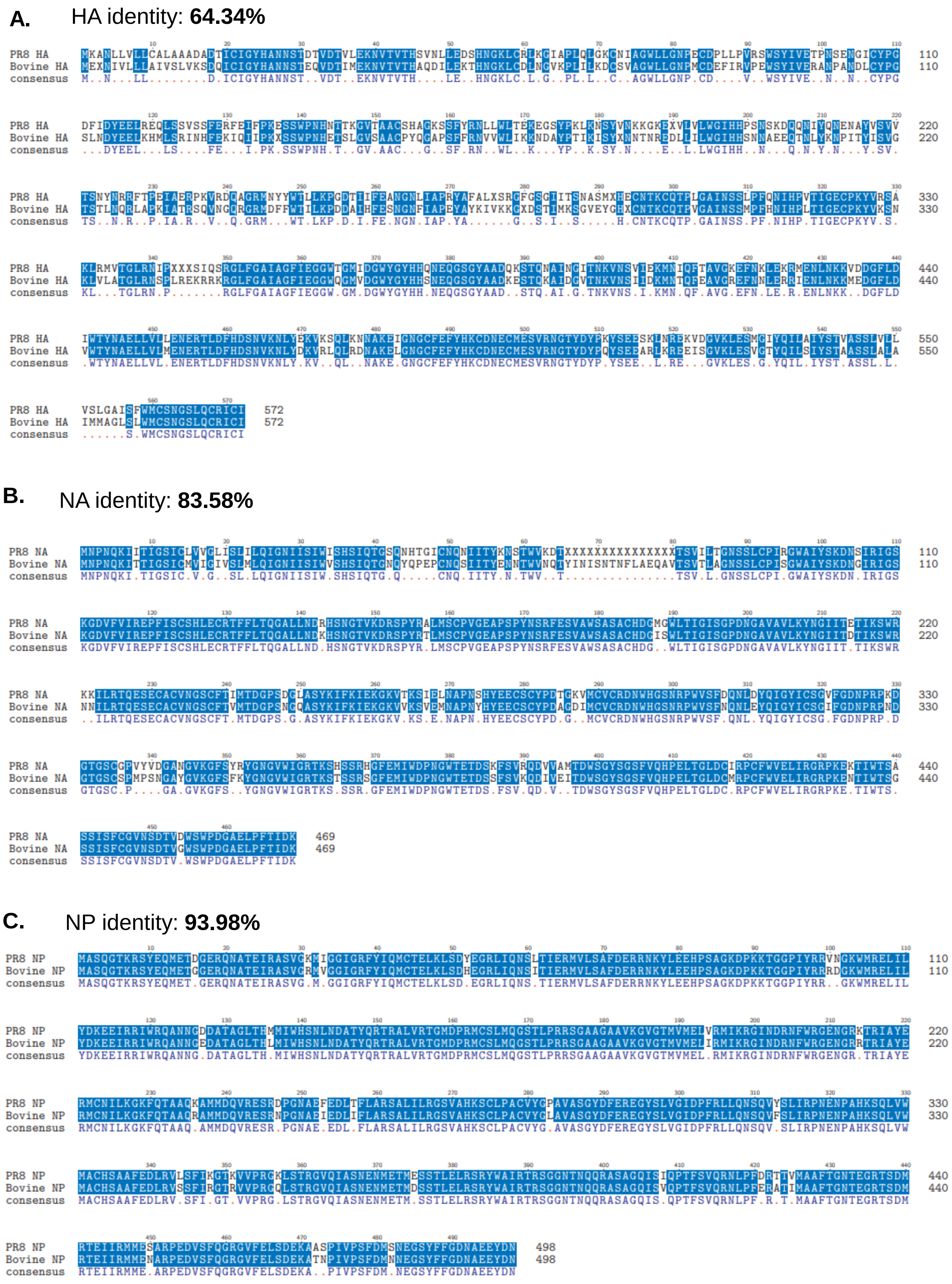


**Supplementary Figure 4.** Comparison of amino acid sequences from H1N1 PR8 and bovine H5N1.
